# Spinal processing of spatiotemporally diverse tactile stimuli: Implications for allodynia and spinal cord stimulation

**DOI:** 10.1101/2025.03.10.642392

**Authors:** Laura Medlock, Steven A. Prescott

**Author notes:** To whom correspondence should be addressed: Steven A. Prescott The Hospital for Sick Children 555 University Avenue Toronto, ON, M5G 1X8 Canada.

## Abstract

Touch is mistakenly perceived as painful when inhibition in the spinal dorsal horn (SDH) is weakened. Disinhibition un-gates a polysynaptic spinal circuit but why mechanical allodynia is predominantly evoked by certain stimuli, like dynamic brushing, remains unclear. To answer this, we incorporated receptive fields (RFs) into a computational model of the SDH to study the processing of stimuli with different spatiotemporal features. Broad stimuli normally suppress output spiking by engaging inhibition from the RF surround, but the efficacy of inhibition depends on the input’s temporal pattern. This is critical since excitatory and inhibitory spinal neurons are preferentially sensitive to synchronous and asynchronous input, respectively. Furthermore, spikes driven by synchronous input are resistant to feedforward inhibition. The combined results explain why broad dynamic touch (e.g. brush or vibration) evokes more allodynia than punctate static touch. Our results also show that asynchronous and spatially disordered input evoked by kilohertz-frequency spinal cord stimulation preferentially activates inhibitory neurons, thus explaining its anti-allodynic effects.

## Introduction

Neuropathic pain patients often experience mechanical allodynia, where touch is perceived as painful^1^. Mechanical allodynia is typically classified as dynamic (evoked by brushing), static (evoked by pressure), or punctate (evoked by von Frey)^2^. These different types of allodynia emerge from touch stimuli with different spatiotemporal features. Static and punctate allodynia both arise from temporally non-fluctuating force; their differences are spatial (broad vs. narrow). Dynamic allodynia arises from force that varies in time (e.g. vibration)^3^ or time and space (e.g. brushing across the skin)^2–4^. Sensing of vibration, which is temporally dynamic, is crucial for the perception of textured surfaces as they are moved across the skin. Texture thus represents a spatiotemporally rich stimulus which causes spatially and temporally patterned activation of primary afferents^5,6^. Such issues have received little attention in the context of allodynia yet dynamic allodynia is common in neuropathic pain^1^, often triggered by clothes moving across the skin.

The spinal dorsal horn (SDH) receives and processes spatiotemporally diverse somatosensory input. This processing relies heavily on synaptic inhibition. Diminished inhibition is one of many changes occurring in the SDH that contribute to abnormal sensory processing in neuropathic pain states^7^. Mechanical allodynia is reproduced experimentally by blocking inhibition, with distinct disinhibitory manipulations yielding similar consequences^8^. Downregulation of the potassium-chloride co-transporter KCC2, which reduces chloride driving force by dysregulating intracellular chloride, is a significant contributor^9^. The molecular signaling and cellular consequences of chloride dysregulation have been thoroughly studied^10^ but the effects on circuit function remain less clear. It is established that disinhibition allows innocuous mechanical input to reach spinal projection neurons (that normally respond exclusively to noxious input) via excitatory interneurons^11,12^. The excitatory interneurons forming this circuitry are necessary for mechanical allodynia^13,14^. Moreover, disinhibition disproportionately affects excitatory interneurons because they rely on strong inhibition to counterbalance strength excitation, whereas inhibitory interneurons balance weak excitation with weak inhibition^15^. Circuit organization clearly influences how disinhibition affects sensation.

Receptive fields (RFs) are valuable for conceptualizing how circuitry affects sensory processing and how (dis)inhibition contributes. Like many neurons, SDH neurons have RFs with an excitatory center and inhibitory surround^15,16^ (**Fig. 1A**). This RF organization reflects the spatial dependence of E-I balance: Stimulation of the RF center evokes net excitation, stimulation of the RF surround evokes net inhibition, and simultaneous stimulation of both regions produces a mixture weighted by the relative input from each area. Because the inhibitory surround is large, broad input engages strong inhibition. That balance shifts towards excitation if inhibition is reduced, encouraging spatial summation^15,16^. However not all inputs are equally affected by inhibition; for instance, temporally synchronous excitatory input can drive rapid depolarization and spiking before inhibitory inputs arrive^17^, producing sensation despite inhibition^18^. Most past studies have not considered the spatiotemporal features of stimuli despite their importance for differentiating allodynia (see above).

**Figure 1.**
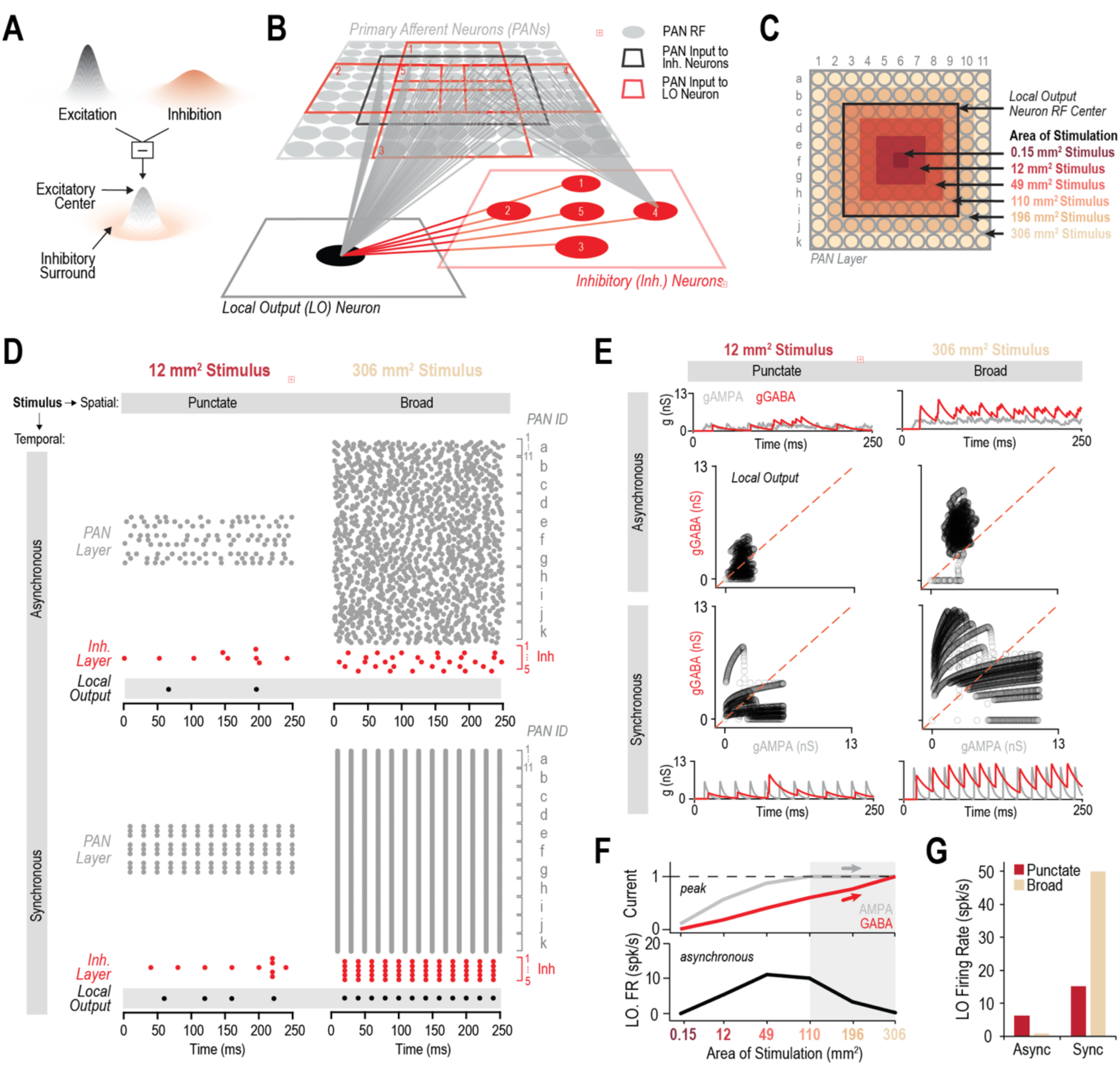
Circuit output depends on spatiotemporal stimulus features. **(A)** Schematic of RF organization. Narrowly distributed excitatory input (gray) and broadly distributed inhibitory input (red) combine to yield an RF with an excitatory center and inhibitory surround. **(B)** Circuitry underlying RF structure. Each PAN has a small RF (gray circle) defined anatomically by its innervation of the skin. local output (LO) neuron RF reflects excitatory input from several PANs plus inhibitory input from inhibitory (Inh.) spinal neurons. Black and red squares denote the RF centers of respective spinal neurons. **(C)** Stimulation area is controlled by varying the number of activated PANs. PAN layer is indexed 1-11 and a-k. **(D)** Raster plot of responses to stimuli with different spatial (punctate, left; broad, right) and temporal (asynchronous, top; synchronous, bottom) features. Each dot represents a spike. **(E)** Inhibitory (gGABA) and excitatory (gAMPA) conductance plots for the LO neuron to stimuli with different spatial and temporal features (same as in D). gGABA (red) and gAMPA (grey) plotted over time or against each other. The latter demonstrate the transient unbalancing of E and I experienced by the LO neuron during synchronous input. **(F)** Normalized current (AMPA, gray; GABA, red) and output neuron firing rate (black) to asynchronous input as stimulus area is increased. Spatial summation leads to increased firing initially (as AMPA current increases) but firing eventually decreases as E-I balance shifts in favour of inhibition (as GABA current continues to increase after AMPA current saturates). **(G)** LO neuron responses vary with stimulus size and temporal pattern. Surround inhibition limits the response to asynchronous input (based on effect explained in **C**) but is ineffective against synchronous input (based on transient unbalancing explained in **B**). LO neuron was delayed-spiking for simulations in **D-F**; inhibitory neurons are always tonic-spiking (see **Fig. 2**).

To investigate how disinhibition disrupts circuit-level processing of somatosensory inputs with different spatiotemporal features, we developed an SDH circuit model that incorporates diverse experimental observations, including RF organization. With this model, we show that surround inhibition limits spatial summation of asynchronous input but not for synchronous input because of the relative timing of excitation and inhibition. Furthermore, the effect of inhibition on spatial summation differed for different inhibitory and excitatory spinal neurons due to differences in their operating modes^19,20^. Consequently, the effect of disinhibition on circuit processing relies on both the temporal patterning of input and the synchrony sensitivity of the spinal neuron receiving it. As such, our model reveals that the processing of synchronous input and its preferential activation of excitatory spinal neurons is disproportionately affected by disinhibition. We also show that effects of disinhibition can cascade within a polysynaptic circuit, leading to a massive expansion of the RF center in downstream neurons, including projection neurons. Lastly, we apply these insights to explain the anti-allodynic effects of kilohertz-frequency spinal cord stimulation (SCS).

## Results

### Inhibition limits spatial summation of asynchronous input but fails to control synchronous input

The center-surround organization of spinal neuron RFs arises from narrowly distributed excitatory input combined with more broadly distributed inhibitory input (**Fig. 1A**). The RF center of a second-order “output” neuron is defined by the cumulative RF of all primary afferent neurons (PANs) converging on it, while its RF surround reflects that cumulative RF of PANs converging on inhibitory interneurons that in turn converge on the output neuron (**Fig. 1B)**. **Figure 1C** illustrates stimulus area relative to PAN RFs. Responses of all cells to asynchronous (e.g. static pressure) or synchronous (e.g. vibration) input of two sizes are illustrated in **Figure 1D**. For asynchronous input, broadening stimulus area from 0.15 to 49 mm^2^ increased AMPA and GABA current in the output neuron, which increased output firing, but further broadening (110 to 306 mm^2^) increased only the GABA current, which reduced output firing (**Fig. 1F**). In other words, spinal excitatory-inhibitory (E-I) balance shifts towards inhibition (gGABA>gAMPA) as the stimulus becomes broader (**Fig. 1E**, top) and thus decreases output firing. However, this effect on output firing was dependent on stimulus timing as broader synchronous input increased, rather than decreased, output firing rates (**Fig. 1G**). This increase occurred because, despite stronger inhibition being engaged (gGABA still >gAMPA), that inhibition arrived on the output neuron too late to prevent spikes driven by synchronous excitatory input (**Fig. 1E**, bottom). These findings demonstrate how spatial and temporal features of the input affect the amount and efficacy of synaptic inhibition, which has major repercussions for SDH circuit output.

### Disinhibition increases spatiotemporal summation in an operating-mode-dependent manner

Given the heterogeneity of SDH neurons and the impact of this heterogeneity on their responsivity to input^20^, we next compared the responses of different types of output neurons to different stimuli under normal and pathological conditions. To asynchronous input (**Fig. 2A**, top), tonic- and delayed-spiking neurons exhibited spatial summation, which was increased by disinhibition, whereas single-spiking neurons did not respond even when disinhibited (due to their narrow integration time window; see **Fig. S7F**). **Figure 2B** reformulates these data for comparison with past experimental data^15^ and confirms that the unmasking of excitation by disinhibition caused a greater increase in firing in excitatory delayed-spiking neurons than in inhibitory tonic-spiking neurons. To synchronous input (**Fig. 2A**, bottom), delayed- and single-spiking neurons were entrained (i.e. responded to each input volley with 1 spike) by stimuli with moderate to large breadth; this was virtually unchanged by disinhibition since feedforward inhibition is already ineffective against synchronous excitatory input (see above). In contrast, disinhibition increased the spiking evoked in tonic-spiking neurons by synchronous inputs because this neuron type relies more heavily on integrating its inputs (see **Figs. S1B** and **S7**). These results confirm that SDH neurons, because of their intrinsic properties (operating mode), integrate inputs differently over time and space, which in turn influences the impact of disinhibition (see **Fig. S2**). While temporal aspects of synaptic integration have been previously considered^21,22^, spatial aspects have received little attention.

**Figure 2.**
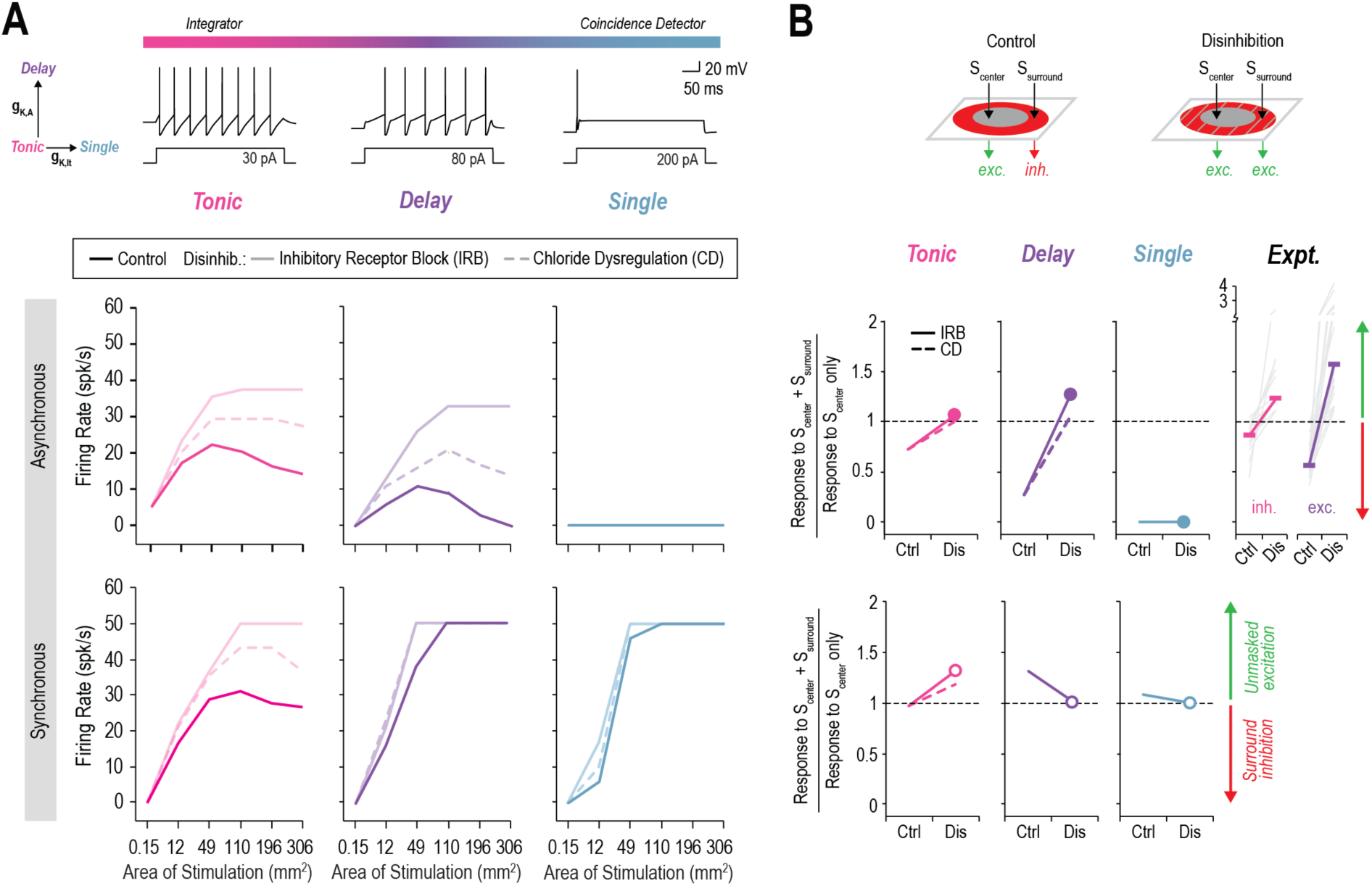
Effects of disinhibition on spatial and temporal summation depend on neuron operating mode. **(A)** Firing rate of tonic- (pink), delayed- (purple), or single- (blue) spiking LO neurons evoked by asynchronous (top) or synchronous (bottom) input of increasing area under control (dark) or disinhibited (pale) conditions. Two forms of disinhibition were simulated (see Methods). **(B)** Ratio of LO neuron responses to stimulation of RF center and surround (S_center_ + S_surround_) compared to stimulation of RF center only (S_center_). The S_center_ was 49 mm^2^ and the S_center_ + S_surround_ was 196 mm^2^ based on experimental data^15^. Stimulation was done under control and disinhibited (IRB, solid; CD, dashed) conditions for asynchronous (top) or synchronous (bottom) input. Similar to experiments (*top right*, Expt), excitatory delayed-spiking neurons respond more to unmasked excitation from the inhibitory surround (ratio >1; green arrow) than inhibitory tonic-spiking neurons following disinhibition.

### Disinhibition has a cascading effect on RF expansion and SDH output

Given the polysynaptic circuitry implicated in allodynia (see Introduction), we explored the effects of RF expansion on downstream SDH neurons by extending the circuit model (**Fig. 3A**) and applying a moving stimulus to map RFs. Second-order local output neurons (blue, orange), which normally respond only weakly to asynchronous input, responded more vigorously when disinhibited, especially for broad input (**Fig. 3B**, top). The third-order global output neuron (black) responded to asynchronous input exclusively when the input was broad *and* the circuit was disinhibited; its RF expanded enough to fill the entire width of the PAN layer, reflecting the compounded effect of RF expansion in upstream local output neurons. This compounding produced even greater RF expansion for higher-order output neurons (**Fig. S3**), which is critical if the polysynaptic circuit upstream of projection neurons includes several excitatory interneurons arranged in series. Synchronous input activated local output neurons more than asynchronous input, where disinhibition only had a modest additional effect (**Fig. 3B, bottom**). The global output neuron only responded to synchronous input following disinhibition, where its RF field expanded, and firing rate increased beyond that evoked by asynchronous input. These results indicate that RF expansion caused by disinhibition can profoundly alter the output of a convergent multi-layer circuit, like in the SDH (see Discussion). Our results also demonstrate that processing of temporally synchronous input (e.g. vibration), and its preferential activation of excitatory spinal neurons, is disproportionately more affected by disinhibition than asynchronous input (**Fig. 3B**). Simulation of dynamic stimuli (brush/texture) further demonstrates that the effect of disinhibition on SDH output also depends on stimulus speed (**Fig. 3C**) and orientation (**Fig. S4**). Consistent with past findings^23–25^, slow brush evoked more global output neuron firing (i.e. more pain) following disinhibition than fast brush because the former activates PANs for longer. PAN activation also depends on stimulus texture, with finer textures evoking faster PAN firing (often ≥200 Hz)^5,26,51,52^, meaning slow brushing with a fine texture causes the greatest output firing (pain).

**Figure 3.**
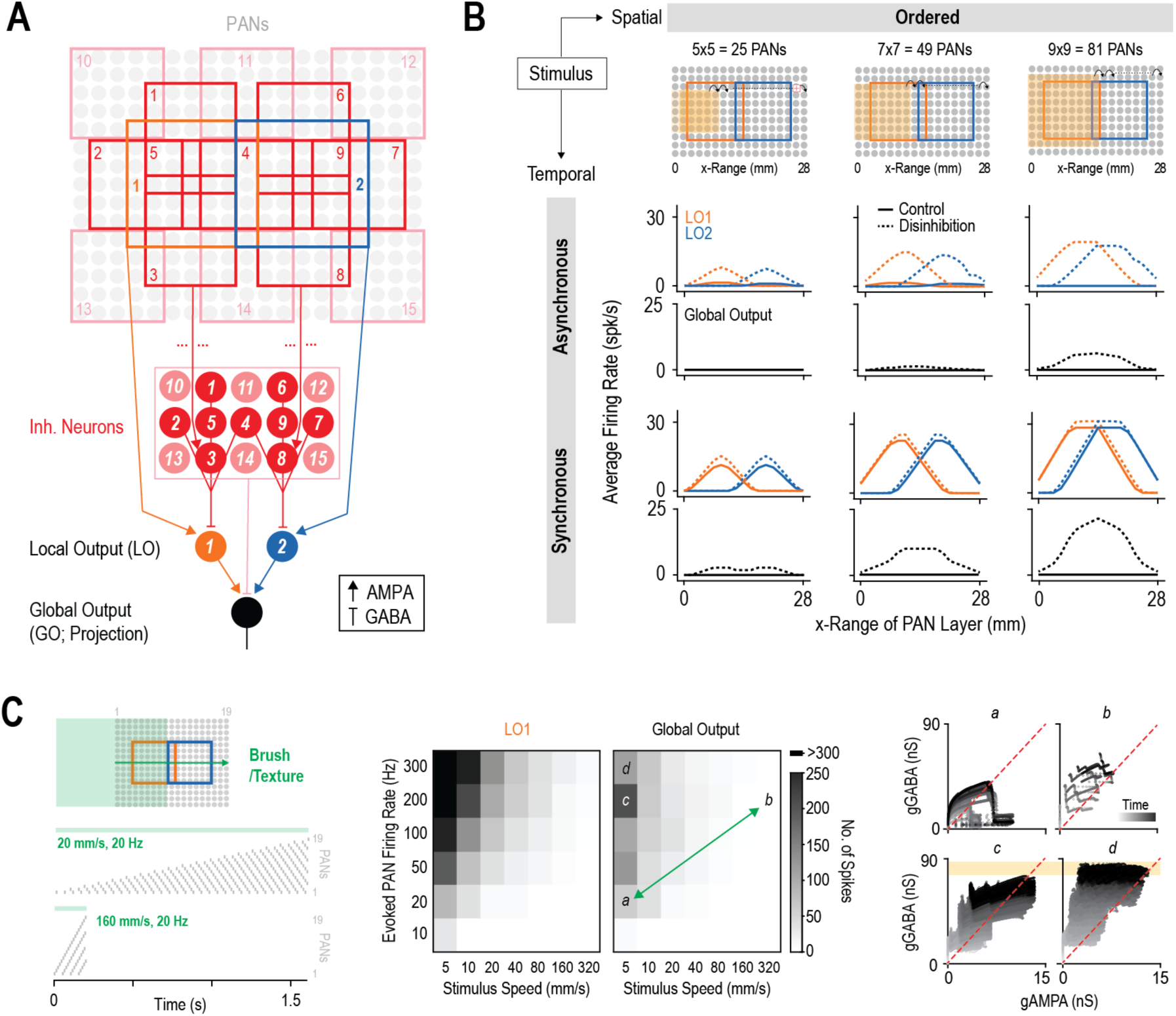
Disinhibition has a cascading effect on RF expansion in the SDH circuit. **(A)** Extended RF circuit. PANs (grey circles), inhibitory neurons (red, pink), local output neurons (LO1, orange; LO2, blue), and a global output neuron (black). LO neurons are both delayed-spiking. Numbered squares outline the RF center for corresponding LO neurons (orange, blue) or inhibitory neurons (red). **(B)** Responses of output neurons to a spatially ordered stimulus (yellow square; moved from 0 to 28 mm in x-Range) with increasing area: (left) 25, (middle) 49, or (right) 81 PANs stimulated. When the input is asynchronous (top) and sufficiently broad (right), disinhibition (IRB) causes RF expansion in LO1 and LO2 (covering ∼66% of PAN layer width) and amplifies RF expansion in the global output neuron (covering the entire PAN layer width). Synchronous input (bottom) has a ceiling effect on RF expansion for LO1 and LO2 but amplifies expansion in the global output neuron (covering the entire PAN layer width). This RF expansion in the global output neuron (and subsequent increase in firing rate) is greater for synchronous input than for asynchronous input. **(C)** Brush simulation. Brush (green, shifting from left to right) causes more intense but shorter-lasting PAN activation when moved slowly (20 mm/s; condition *a*) than quickly (160 mm/s; condition *b*) (see rasters on left). Center panels summarize activation of local and global output neurons as a function of brush speed and PAN firing rate, where the latter depends on brush speed and texture (see text) following disinhibition (CD, E_GABA_ to -50 mV). The Global Output neuron was not activated by any stimuli under control conditions. For the same texture brushed at different speeds, slow brushing (*a*) evoked more output firing than fast brushing (*b*). For the same slow brush speed, fine textures (*c* and *d*) evoked more output firing than a coarse texture (*a*). Right panels compare E-I balance for stimulus conditions *a-d*. Inhibition (gGABA) accumulated quicker than excitation (gAMPA) during fast brushing (*b*), evidenced by data points staying above the diagonal, whereas excitation exceeded inhibition more often during slower brushing (*a*) as revealed by excursions to the right of the diagonal. There is a ceiling effect on LO1 firing rates from *c* to *d*, but the additional recruitment of inhibition (# of Inh. Neurons) causes a decrease in Global Output firing in *d*.

### Spatiotemporally disorganized input preferentially engages spinal inhibition and combats pathological disinhibition

Until now, all inputs were treated as spatially continuous to simulate tactile stimuli uniformly activating PANs with contiguous RFs (see **Fig. 1**). PANs can be activated in a discontinuous or patchy manner by patterned stimuli (e.g. different textures) but the resulting input is complex^26^ and has not been thoroughly studied in rodents. A random or disorganized pattern of PAN activation also arises from extracellular electrical stimulation of peripheral nerves or dorsal columns (e.g. during SCS) because PAN axons are not somatotopically organized to the spatial resolution of individual RFs; instead, axon size and position within the electric field dictate which axons are recruited, with more intense pulses activating progressively smaller axons and those farther from the stimulating electrodes^27^. Intensity is parameterized as the fraction of PANs recruited (see **Fig. 4A**). Conventional, low-rate (50 Hz) stimulation activates PANs synchronously whereas high-rate (1 kHz) stimulation activates PANs asynchronously but at roughly the same firing rate because of differences in entrainment (**Fig. S5**)^18^.

**Figure 4.**
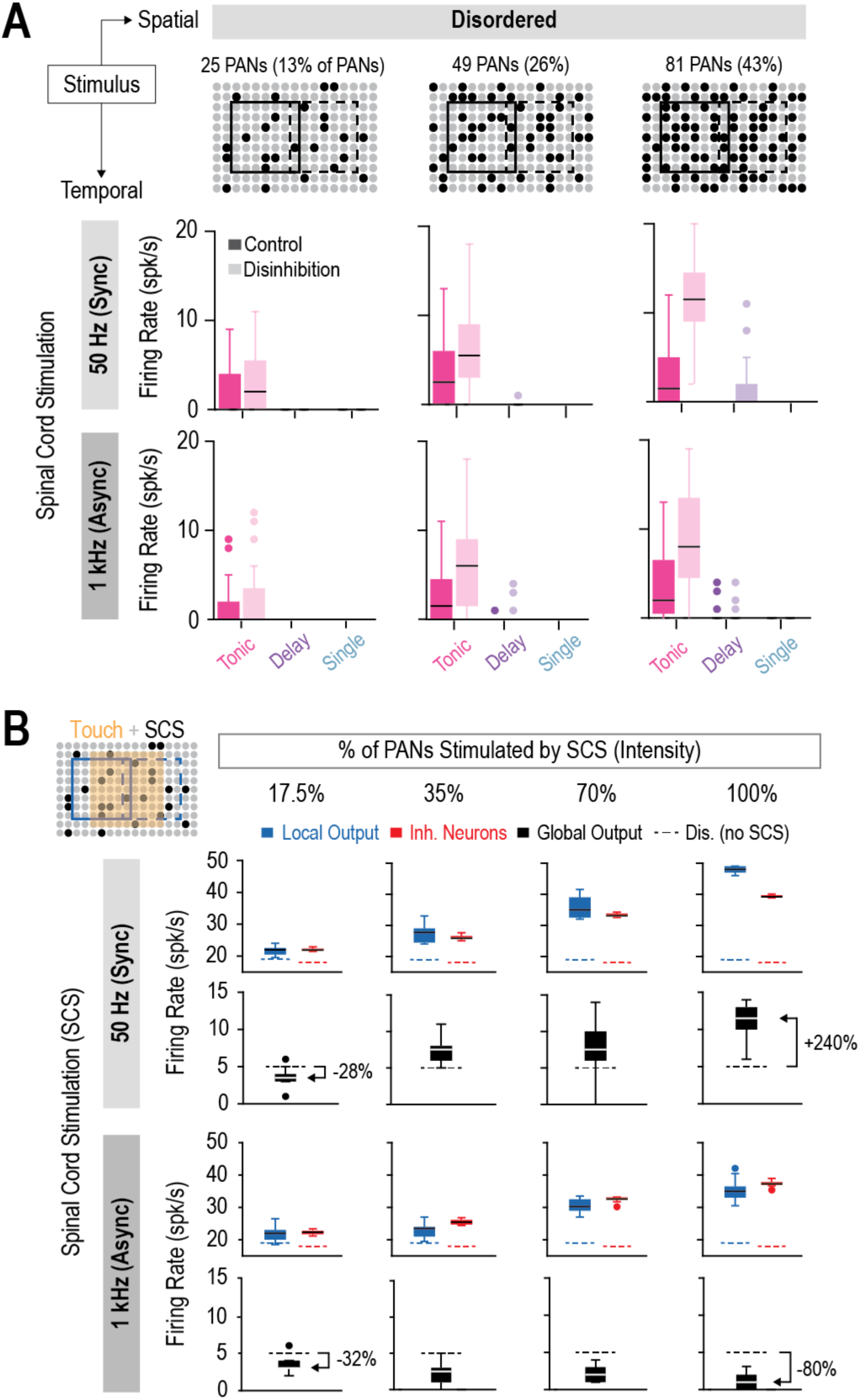
Spatiotemporally disorganized input preferentially engages spinal inhibition and combats the effects of disinhibition. **(A)** Responses of tonic- (pink), delayed- (purple), or single- (blue) spiking neurons to spike trains recorded from PANs during 50 Hz (top) or 1 kHz (bottom) SCS)^18^ under control (dark) or disinhibited (pale, IRB) conditions. SCS evokes spatially disordered input (black dots = stimulated PANs) that is synchronous (50 Hz SCS) or asynchronous (1 kHz SCS). To match **Figure 3B**, either 25, 49, 81 PANs were activated (left to right) to simulate increasing SCS pulse intensity. Activated PANs were randomly selected per trial (n=20 trials/stimulus type). Both SCS types preferentially activated tonic-spiking (inhibitory) neurons. **(B)** Modulation of touch-evoked responses by SCS. Spatially ordered touch input (beige shading) and disordered SCS input (black dots) were applied to extended SDH circuit (from **Fig. 3A**). Dashed lines show touch-evoked responses after disinhibition (E_GABA_ to -55 mV); output neurons did not respond to touch alone under control conditions. SCS increased firing in LO (blue) and inhibitory (red) neurons, which, depending on their relative activation, increased or decreased firing in global output (black) neurons. All output neurons are delayed-spiking. Whereas 50 Hz SCS (top) preferentially activated inhibitory neurons only for weak pulses (left), 1 kHz SCS (bottom) preferentially activated them across all pulse intensities, causing a robust reduction in global output neuron firing.

**Figure 4A** shows that spatially disordered input preferentially activated tonic-spiking neurons (pink), with delay- (purple) and single-spiking (blue) neurons responding weakly or not at all. This was true for both 50 Hz and 1 kHz stimulation, and across a range of pulse intensities. This differential activation occurs because delayed- and single-spiking neurons need relatively strong excitatory input, which requires concentrated activation of their RF center (which is unlikely to occur with spatially disorganized PAN activation), whereas tonic-spiking neurons can be activated by weaker input because they integrate it over time (**Fig. S7**). Since most inhibitory neurons are tonic-spiking whereas delayed- and single-spiking neurons are mostly excitatory, these results mean SCS preferentially activates inhibitory SDH neurons. This is taken for granted since SCS reduces pain, but a mechanistic explanation has been elusive since the PANs activated by SCS provide synaptic input to both inhibitory *and* excitatory SDH interneurons; indeed, the latter are presumably activated when SCS evokes uncomfortable paresthesia.

To explore how SCS reduces allodynia, we used the SDH circuit in Figure 3 to simulate responses to touch *and* SCS under disinhibited conditions (Fig. 4B). The firing rates evoked by touch input alone are plotted as dashed lines to highlight changes in firing due to SCS applied at four different intensities. For 50 Hz SCS, weak pulses activated excitatory (local output, blue) and inhibitory interneurons (red) with the net effect of reducing activation of the global output neuron (black); but as pulse intensity was increased, local output neurons were more strongly activated by synchronous 50 Hz SCS input than inhibitory neurons (see Fig. 2), resulting in increased activation of the global output neuron, which would increase pain. Preferential activation of excitatory (delayed-spiking) neurons by SCS seems to contradict results in Figure 4A, but whereas panel A considers the response to SCS-evoked input by itself, panel B considers the combination of SCS- and touch-evoked input. Because excitatory neurons prefer synchronous input (see Fig. 2), they respond well to input evoked by 50 Hz SCS when it is superimposed on asynchronous touch input, even if the former is spatially disordered. These results are consistent with 50 Hz SCS producing analgesia over a relatively narrow range of pulse intensities before uncomfortable paresthesia develops with stronger pulses. By comparison, during 1 kHz SCS, inhibitory interneurons were more strongly activated than local output neurons across the full range of pulse intensities, resulting in reduced activation of the global output neuron, which should robustly relieve pain. This is consistent with inhibitory (tonic spiking) neurons preferring asynchronous input, even when it is spatially disordered. Together, these results reveal that spinal E-I balance depends on the spatiotemporal features of the stimulus and that spatiotemporally disorganized input preferentially engages spinal inhibition and can combat the effects of pathological disinhibition.

## Discussion

This study used computational modelling of the SDH circuit to examine how somatosensory inputs with different spatiotemporal patterns are normally processed, and how pathologically altered processing leads to allodynia. Our results show that a broad stimulus, by extending into the inhibitory RF surround, engages more inhibition than a narrow stimulus, but the effect on output neuron spiking depends on the input’s temporal pattern (Fig. 1). The differential sensitivity of excitatory and inhibitory spinal neurons to input synchrony exacerbates the effect of stimulus temporal patterning (Fig. 2). Effects of disinhibition also depend on a neuron’s position in the circuit, with downstream neurons experiencing greater RF expansion than upstream neurons (Fig. 3). Our results also show that kHz-frequency SCS produces spatiotemporally disorganized input that suppresses touch-evoked spiking in output neurons by preferentially activating inhibitory neurons without producing synchronous excitation that might activate excitatory neurons (Fig. 4). These results highlight how cell and circuit properties affect SDH function, including why brushing – a broad stimulus causing synchronous inputs (Fig. 3C) – is especially prone to causing allodynia^23,24^. Beyond explaining clinical symptoms, such information can help optimize therapeutic interventions.

Different types of allodynia are associated with different pain etiologies and likely arise through different biological mechanisms^2,28^. For instance, touch signals could be relayed centrally by sensitized nociceptors, or they could be relayed normally (by low-threshold mechanoreceptors) but get misprocessed in the SDH. The latter occurs in dynamic allodynia, but diverse pathological changes are not mutually exclusive and may instead combine. Using clinical phenotype to infer underlying biological changes has been repeatedly espoused^29–32^, but the time has come to replace “cartoon” models with quantitative ones. Mechanistic models, like ours, can help interpret past observations. For instance, Samuelsson et al.^33^ showed that dynamic allodynia scales with the number and length of brush strokes (as expected) but not with brush width (contrary to expectations), but this likely depends on brush width *relative to RF size* (**Fig. S6**).

Tuning curves (which include RFs) describe the dependence of a neuron’s response on stimulus location, intensity, or other features. Neural tuning affects how information is represented^34^. Deciphering the biological basis for neural tuning (e.g. Angelucci et al.^35^) thus advances our understanding of sensory processing, especially when combined with mechanistic modelling (e.g. Di Santo et al.^36^). Neural coding in the visual system is much better understood than in the pain system, but some tentative comparisons are useful. Stimulation outside a visual neuron’s excitatory RF center normally suppresses firing, but not always; for instance, neurons in a fluctuation-driven regime (as opposed to mean-driven) may fire more^37^. This is consistent with our results: Stimulation in the inhibitory RF surround suppressed spiking driven by asynchronous input (i.e. in a mean-driven regime) but enhanced the spiking driven by synchronous input (i.e. in a fluctuation-driven regime) (see Fig. 1F). Stimulation in the inhibitory RF surround causes a net increase in inhibition, but the effect on spiking depends on the relative timing of excitation and inhibition^17,18,38^. Two factors are important. First, inhibition driven by synchronous input often arrives at the output neuron too late to prevent the rapid depolarization evoked by synchronous excitation from triggering a spike. Second, stimulation in the RF surround engages some excitation (albeit less than inhibition), but whereas inhibition arrives late, excitation from the RF surround arrives at the same time as excitation from the RF center, summating so that otherwise subthreshold voltage fluctuations reach threshold. Just as spatiotemporal context helps explain visual illusions and after-effects^39^, understanding of allodynia may likewise benefit.

Ideally one can harness a deeper mechanistic understanding of allodynia to devise better strategies to alleviate it. Beyond restoring chloride regulation^40^ or otherwise strengthening inhibition pharmacologically, one should consider complementary interventions. Indeed, if SCS acts by engaging inhibition^41^, then its ability to reduce mechanical allodynia^42^ implies residual inhibition can be engaged, and that doing so reduces output neuron spiking. But how is it that activating low-threshold mechanoreceptors with SCS does not itself cause allodynia? Our results suggest that SCS preferentially activates inhibitory neurons because it triggers spatially disorganized input, meaning a neuron’s input arrives mostly from its inhibitory surround without any spatially concentrated input from its excitatory center. Furthermore, during kilohertz-frequency SCS, the input desynchronizes^18^, becoming temporally disorganized, which also favors activation of inhibitory neurons. Gilbert et al.^43,44^ invoked lateral inhibition to help explain the effects of SCS, but they did not formally model center-surround RF organization. Another critical difference is that excitatory interneurons received excitatory input exclusively from high-threshold C fibres^44^, contrary to known connectivity^14^ and the necessity of excitatory interneurons for mechanical allodynia (see Introduction). Notably, the absence of input from low-threshold fiber precludes SCS from activating excitatory interneurons.

To summarize, our results demonstrate how the balance of excitation and inhibition varies over space and time depending on (1) the spatiotemporal features of the input and (2) the circuit and constituent neurons processing that input. This is a more sophisticated explanation of E-I balance than is typically considered, but such details are necessary to more fully explain allodynia and to more effectively treat it.

## Materials & Methods

### SPINAL NEURON MODELS

SDH neurons are heterogeneous. To reflect that heterogeneity, we simulated neurons with the most common spiking patterns, namely integrators (tonic-spiking), coincidence detectors (single-spiking), and a hybrid of these operating modes (delayed-spiking) (**Fig. S7**). Based on experimental findings^45–47^, inhibitory neurons are typically tonic-spiking whereas excitatory neurons are either delayed- or single-spiking. Spinal neurons were simulated as single-compartment adaptive exponential (AdEx) models^48^ with parameters: a = 0 nA, b = 0 nA, Δ_T_ = 2 mV, V_T_ = -45 mV, t_ref_ = 5 ms, V_reset_ = -85 mV (or -70 mV for single spike cells). To the AdEx models, we added a leak current (I_leak_), as well as fast (I_adapt-fast_) and slow (I_adapt-slow_) adapting currents^20^, and additional currents based on neuron type: Tonic-spiking neurons had a subthreshold inward current (I_sub_)^20^, delayed-spiking neurons had an A-type K^+^ current (I_K,A_)^45^ and single-spiking neurons had a low-threshold K^+^ current (I_K,lt_)^45^ (**Fig. S7A**). Equations for these currents are described in **Supplemental Table 1**. Parameters for additional currents were as previously published^20,45^, except for τ_adapt-slow_ = 40 ms, τ_adapt-fast_ = 5 ms. Maximal conductance densities (g_max_) and other details are summarized in **Supplemental Table 2**. Reversal potentials were E_Na_ = 50 mV, and E_K_ = -100 mV, E_leak_ = -70 mV for all cells. Membrane capacitance (*C_M_*) is 1 μF/cm^2^ for all neurons. The spike-triggered average (STA) for different neurons was calculated for each neuron type by injecting either fast (τ_noise_=5 ms) or slow (τ_noise_=100 ms) noise simulated as an Ornstein–Uhlenbeck (OU) processes as previously described^20^ (μ_delay_=0.05 nA, μ_tonic_=0.003 nA, μ_single_=0.22 nA; σ_all_ = 1.2 nA) into the soma (**Fig. S7F**).

### SDH CIRCUIT MODEL

#### NEURON POPULATIONS & INPUT

Spinal neuron RFs have an excitatory center and inhibitory surround (Fig. 1A). In the model, this structure is formed via excitatory input from an 11×11 grid of primary afferent neurons (PANs, grey) and inhibitory input from 5 spinal neurons (Inh. Neurons 1-5, red) which connects to 1 local output (LO) spinal neuron (black) (Fig. 1B). The excitatory center of the LO neuron is formed by a 7×7 grid of PANs (PAN→LO) and the inhibitory surround is formed by input from I_1_-I_5_ neurons (Inh.Neurons→LO). Inhibitory neurons also receive excitatory input from PANs (PAN→Inh.Neurons; n=25 synapses/neuron) but receive no inhibitory input. The RFs for I_1_-I_5_ (red squares) are tiled on and around the excitatory RF center (black square) of the LO neuron and constitute its inhibitory RF surround (Fig. 1B). The LO neuron was modeled with different spiking patterns (see *Spinal Neuron Models*) for comparison, but for more complex circuit models (Figures 3-4), the LO neuron is treated as excitatory and modeled as delayed-spiking. Inhibitory neurons I_1_-I_5_ are modelled as tonic-spiking. Touch input (1 s long) to the PANs was simulated as a NetStim of 50 Hz with different noise parameters for asynchronous (noise = 0.6) or synchronous (noise = 0) input (**Fig. S8A**). Each PAN was simulated as a separate NetStim to allow for heterogeneous input. Spinal cord stimulation (SCS) input was simulated by injecting PANs with a VecStim of different experimental spike trains recorded from primary afferents during either 50 Hz or 1 kHz SCS^18^ (**Fig. S5**).

#### SYNAPTIC CONNECTIVITY

Synaptic connectivity is shown in **Figure S8A**. Excitatory synaptic transmission was mediated by AMPA receptors, whereas GABA receptors mediated inhibitory synaptic transmission. Synapses were modelled using the Exp2Syn mechanism with rise and decay time constants (AMPA: 0.1 ms, 5 ms; GABA: 0.1 ms, 20 ms) and scaled by a synaptic weight (described below). *E*_GABA_ was -70 mV and *E*_AMPA_ was 0 mV. Pathological disinhibition was simulated as complete inhibitory receptor block (IRB) by setting g_GABA_=0 or as chloride dysregulation (CD) by shifting *E*_GABA_ from -70 to - 55 mV (or -60 mV for a subtler effect in Figure 2).

#### SPATIAL ORGANIZATION

Spinal neuron RFs are defined by the summation of spatially distributed input. Specifically, narrow excitation (grey dashed) combined with broad inhibition (red dashed) produces a net RF with an excitatory center and inhibitory surround (black; **Fig. S8B**). RF architecture was based on size and firing rate of RFs from experimental data^15,49^. Specifically, the inhibitory surround is larger than the average excitatory center (∼50 mm^2^) for lamina I/II spinal neurons^15^. To account for the spatial distribution of input, PANs and inhibitory neurons have a 2D spatial organization (see Fig. 1B) and the synaptic weights were spatially dispersed according to this organization (**Fig. S8C**).

The entire layer of PANs is 17.5 x 17.5 mm with 1.75 mm grid spacing and centered at (8.75 mm, 8.75 mm) in (x,y) space. The layer of Inh. Neurons is 3.5 x 3.5 mm with the same grid spacing and (x,y) centering. Excitatory and inhibitory weights to the LO neuron were dispersed over the x-y plane of the PAN and inhibitory neuron layers, respectively (**Fig. S8C**). This ensured that weights were proportional to the distance of the presynaptic cell’s RF from the center of the postsynaptic cell’s RF (i.e. that weights were smaller if RFs were farther apart). Experiments show that excitatory spinal neurons receive strong excitation and inhibition, whereas inhibitory neurons receive weaker excitation and inhibition^15^, which contributes to their differential net RF (**Fig. S8B**). To account for these differences, synaptic weights (*w_syn_*) were scaled by *w_max_* based on the LO neuron type (Exc. LO or Inh. LO). Specifically, PAN→Inh.Neurons, Inh.Neurons→InhLO, and Inh.Neurons→ExcLO weights were dispersed using a Gaussian function,

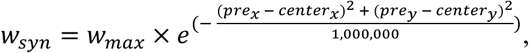

where *w_max_* is the maximum synaptic weight at the RF center, *pre_x,y_* is the x- or y-coordinate of the presynaptic cell, and *center_x,y_* is the x- or y-coordinate of (1) the postsynaptic cell’s RF center for AMPA synapses or (2) the inhibitory neurons whose RF completely overlaps the LO neuron’s RF (e.g. Inh_5_ for the single RF circuit) for GABA synapses. Based on experimental findings^15^, the spatial dispersion of PAN→ExcLO weights was described by the sum of two Gaussians,

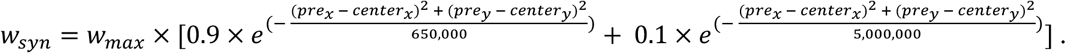

The *w_max_* for AMPA and GABA synapses were 0.001 and 0.002 for Exc. LO neurons and 0.0005 and 0.001 (i.e. 2x smaller) for Inh. LO neurons. Dispersion of PAN→Inh.Neurons weights was the same as PAN→InhLO. Simulations were conducted in NetPyNE ^50^.

#### EXTENDED SDH CIRCUIT

The extended RF circuit (Fig. 3A) contains two LO neurons (LO1, orange, LO2, blue) and a global output neuron (GO, black). The layer of PANs was extended to 771.75 mm^2^ (n=285 neurons) and the layer of inhibitory neurons extended to 24.5 mm^2^ (n=15 neurons). LO1 received inhibition from Inh_1_-Inh_5_ neurons and LO2 from Inh_4_, and Inh_6_-Inh_9_. The global output neuron received broader inhibition from all Inh_1_-Inh_15_. The circuit connectivity is shown in Figure 3A. The rules of spatial dispersion for LO1 and LO2 remain as previously described. The AMPA and GABA synaptic weights to the global output neuron are 0.005 and 0.0025.

### CODE AVAILABILITY

The model code is available here: https://modeldb.science/2018015

## Supplemental Figures

**Figure S1.**
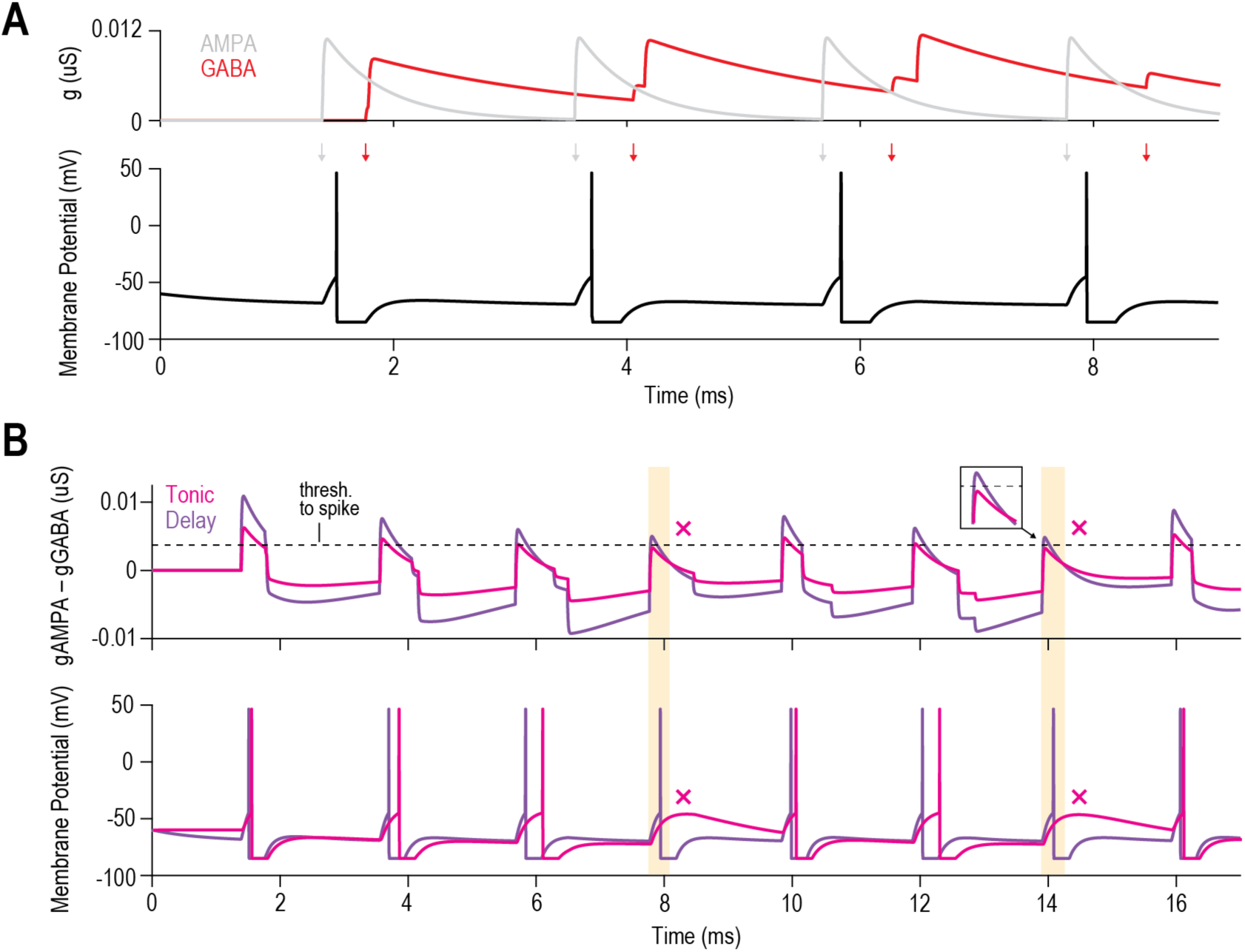
Influence of input synchrony on output neuron firing (extension of Figure 1). (A) Feedforward inhibition (GABA, red) arrives too late to prevent the output neuron from responding to synchronous excitatory input (AMPA, grey). (B) When input is synchronous, tonic-spiking neurons (pink) are more affected by inhibition than other spinal neuron types. As integrators (see Fig. S7), tonic-spiking neurons summate input more slowly than other spinal neurons (e.g. delayed-spiking, purple) and receive 2x weaker input (g_AMPA_ and g_GABA_) based on experimental findings^18^. For a neuron to reach spike threshold at the arrival of AMPA input, gAMPA must exceed gGABA by a critical amount (dashed line). If enough gGABA is left over from the previous cycle (yellow boxes), the tonic-firing neuron fails to reach threshold (pink x’s). For delayed- and single-spiking neurons, g_AMPA_ consistently exceeds g_GABA_ when input arrives due to their larger weights (see *Methods*), so that a spike occurs consistently on each cycle.

**Figure S2.**
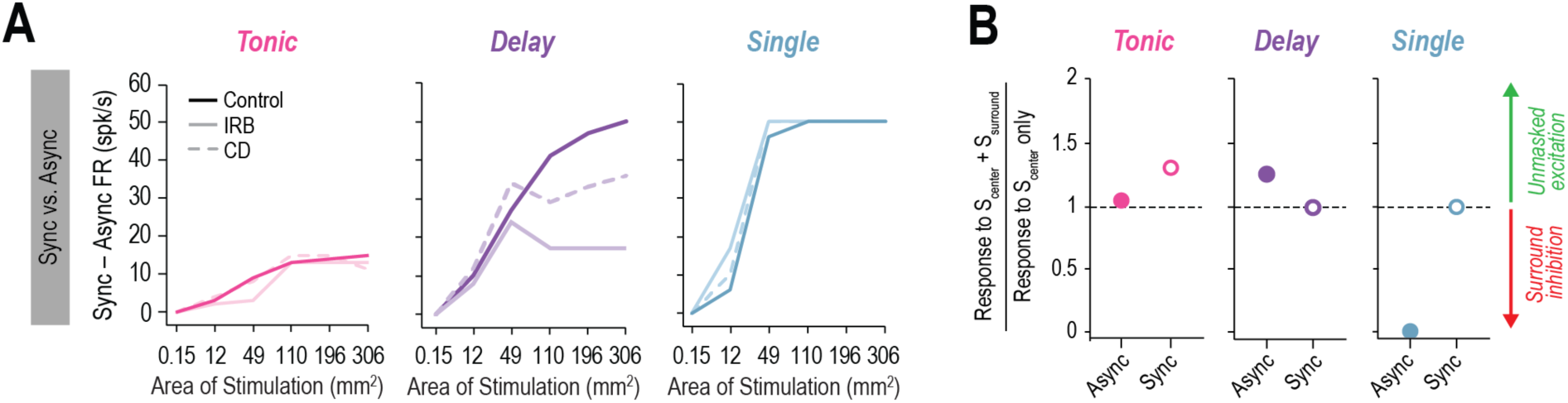
Preference for synchronous input differs across spinal neuron types (extension of **Figure 2**). **(A)** Difference in firing evoked by synchronous and asynchronous input for equivalent area. Single-spiking neurons strongly prefer synchronous input, as do delayed-spiking neurons, although less so when disinhibited. Tonic-spiking neurons exhibit a weak preference for synchronous input. **(B)** Ratio of firing evoked by co-stimulation of RF center and surround (S_center_ + S_surround_) to stimulation of RF center only (S_center_) when disinhibited. Ratio >1 demonstrates excitation unmasked by reducing surround inhibition. For tonic-spiking neurons, unmasking synchronous input had a stronger effect than unmasking asynchronous input, and vice versa for delayed-spiking neurons.

**Figure S3.**
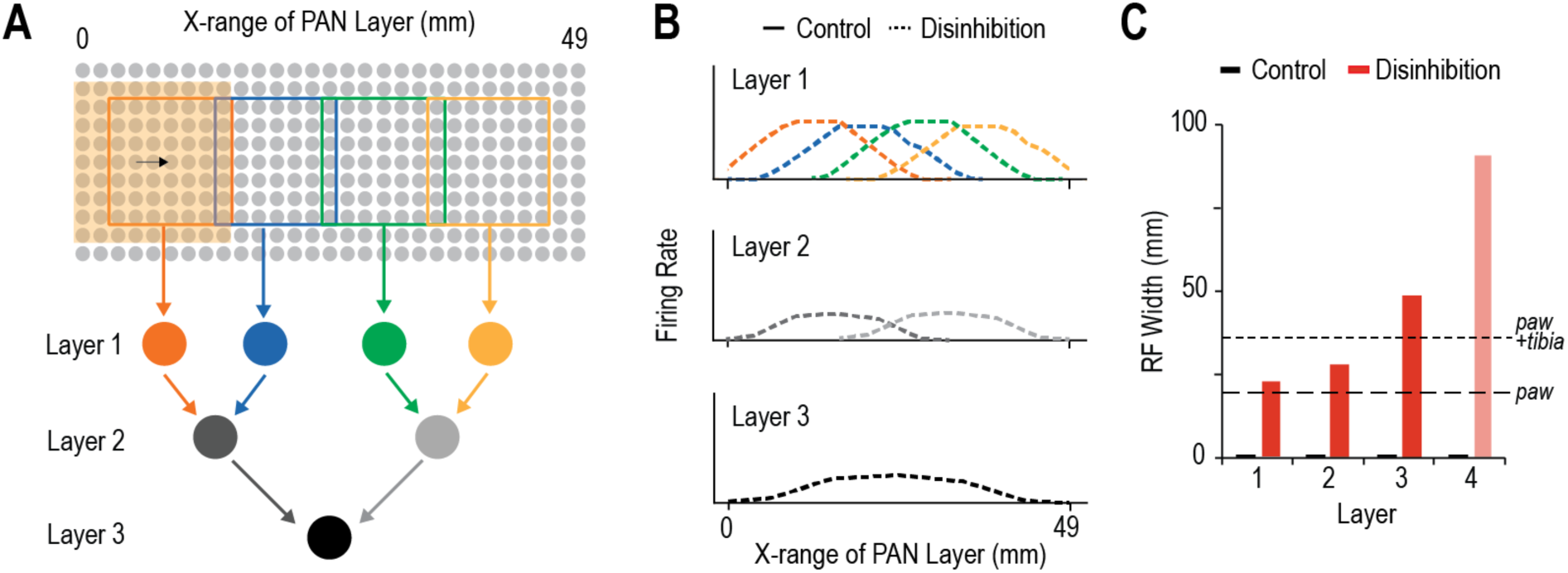
Disinhibitory effects cascade, causing progressive expansion of RFs in higher-order neurons (extension of **Figure 3**). **(A)** Schematic of a polysynaptic circuit with three converging layers. Output neurons are depicted; inhibitory neurons are not shown. **(B)** Disinhibition widens the RF and increases firing evoked by asynchronous input in layer 1, 2 and 3 output neurons allowing those neurons to now respond to activation of ∼50% (24.5 mm), ∼66% (33 mm), or 100% (49 mm) of the PAN layer, respectively. All output neurons are delayed-spiking. Layer 2 and 3 neurons do not respond to asynchronous input under normal (control) conditions. **(C)** The RF expansion due to disinhibition (red) increases exponentially for neurons farther downstream. Length of a mouse paw^53^ (long dash) and mouse paw+tibia^54^ (short dash) are shown for comparison.

**Figure S4.**
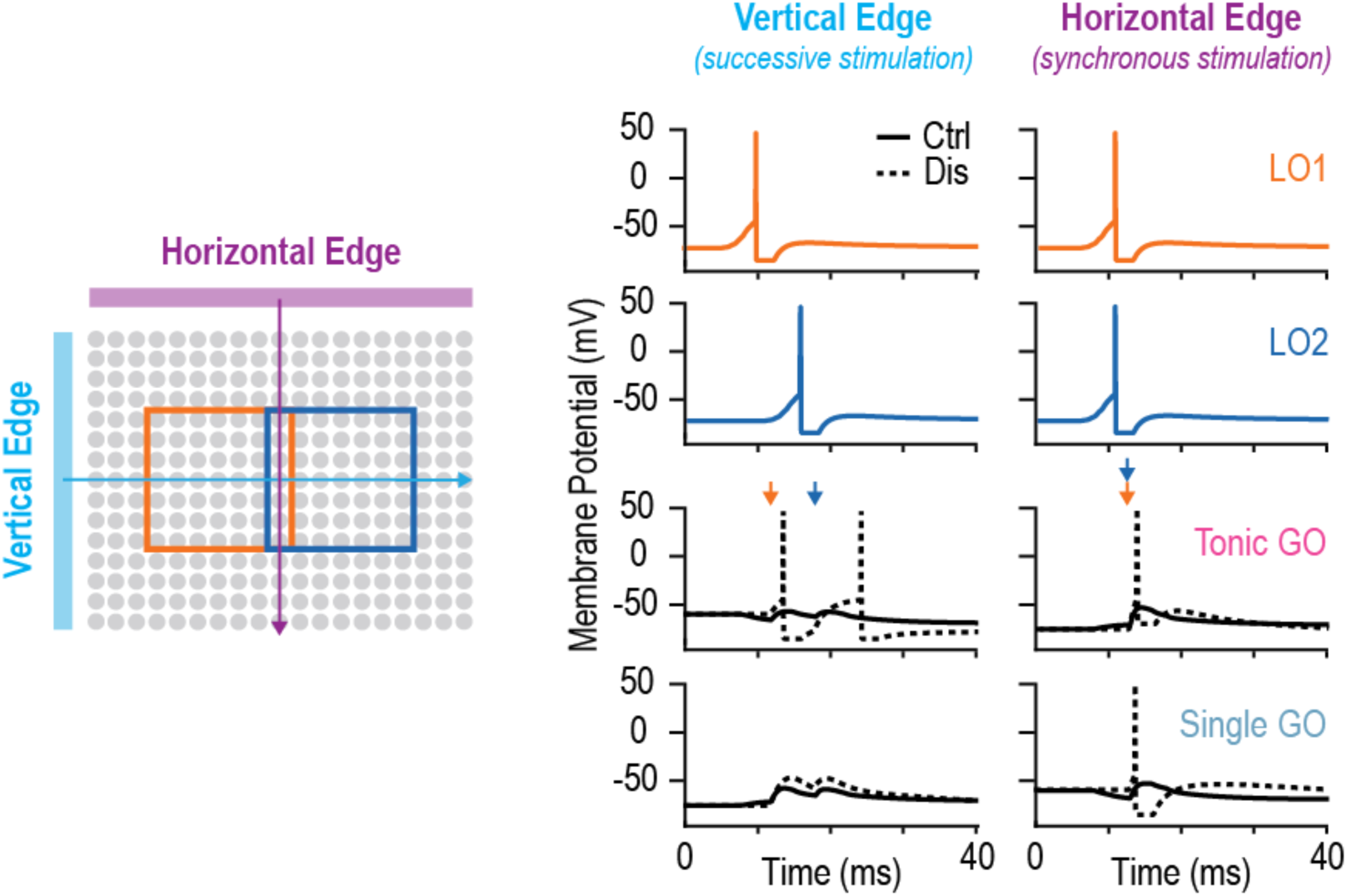
Stimulus orientation affects SDH output following disinhibition (extension of **Figure 3**). (left) Schematic of a vertical (blue) or horizontal (purple) edge that results in successive or synchronous activation, respectively, of the two LO neurons from the network shown in Figure 3A. The vertical edge first activates LO1 followed by LO2 a few milliseconds later (asynchronously) whereas the horizontal edge activates LO1 and LO2 at the same time (synchronously). Under normal conditions, neither stimulus orientation activates the Global Output (GO) spinal neuron, regardless of GO neuron operating mode (tonic-spiking, pink; single-spiking, blue). However, following disinhibition (IRB, dashed lines), the tonic-spiking (integrator) GO neuron spikes because it summates the excitatory inputs over time. The single-spiking (coincidence detector) GO neuron does not respond because it cannot summate input (see Fig. S7). Conversely, synchronous stimulation of LO1 and LO2 by a horizontal edge excites both tonic- and single-spiking GO neurons following disinhibition because there is no need for temporal summation. Here the speed of the stimulus across the PAN layer is 1.5 m/s to allow for quick temporal summation and spiking in LO1 and LO2 and is similar to previous experiments^25^.

**Figure S5.**
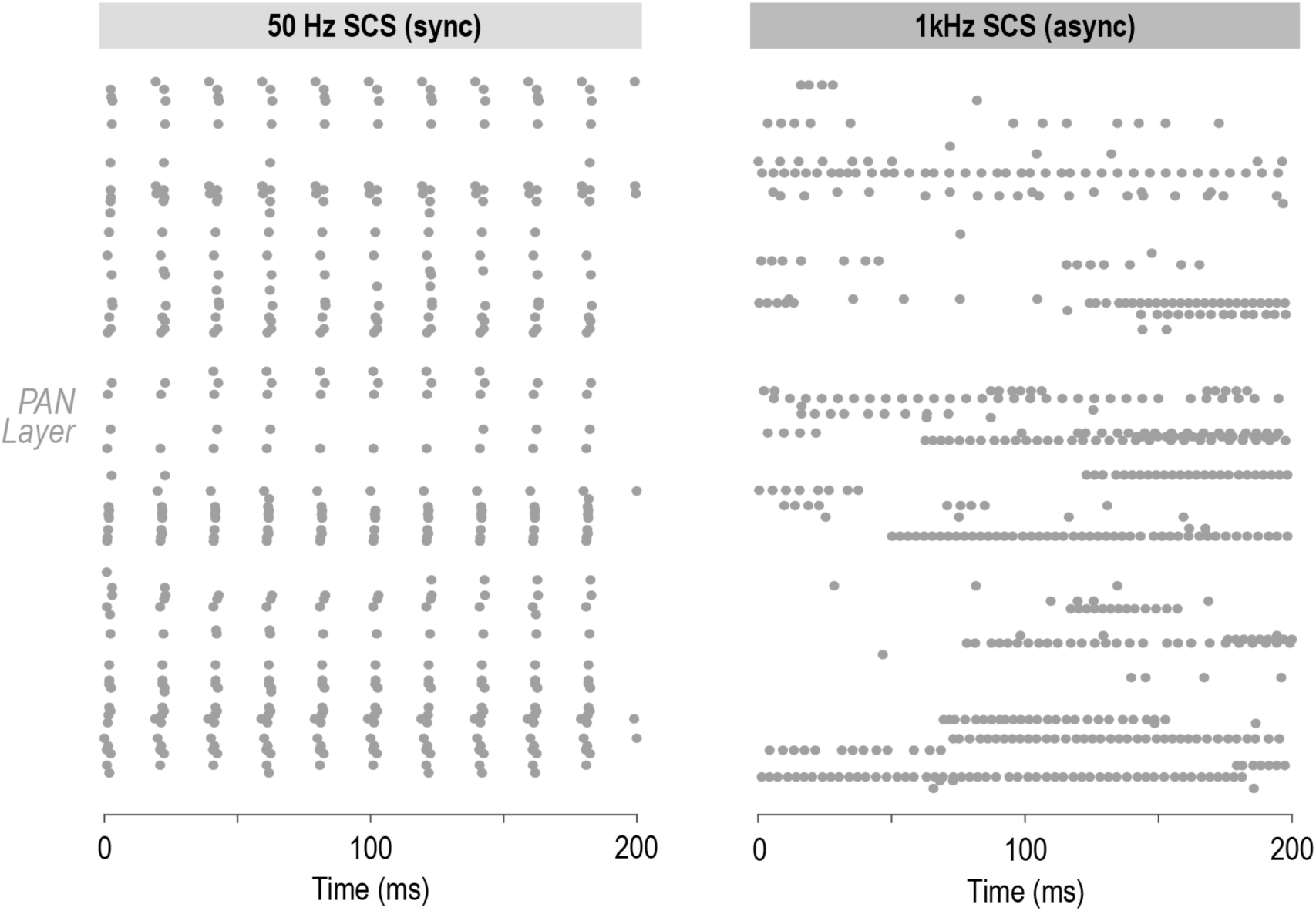
Raster plot of PAN responses to 50 Hz (left) and 1 kHz (right) SCS recorded from urethane-anesthetized rates (from ref.^18^). PANs respond synchronously to 50 Hz SCS and asynchronously to 1 kHz SCS.

**Figure S6.**
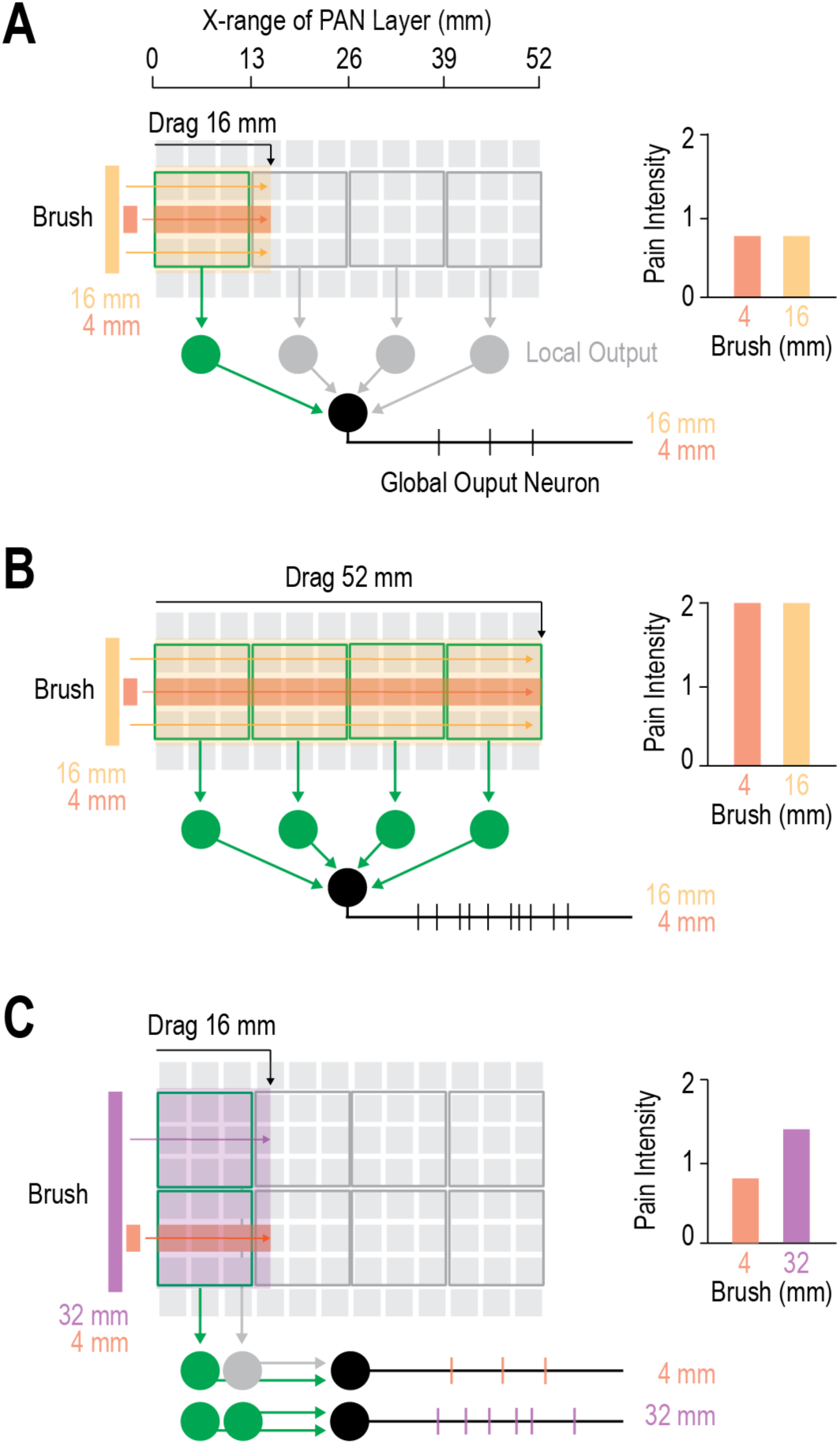
Impact of stimulus design on experimental or clinical probing of mechanical pain. Light grey squares represent PAN RFs; larger boxes represent LO neuron RFs. Global Output neuron is shown in black. RF sizes are scaled to data from the glabrous skin of the human hand (palm): PAN RF = 16 mm^2^ (4 mm x 4 mm)^55,56^, SDH RF = 169 mm^2^ (13 mm x 13mm) [calculated based on the ratio of human (hand) to rat (hindpaw) PAN RF size (ratio = 16 mm^2^ / 4 mm^2^ [57, 58] = 4)]. Spinal RFs in the rat hind paw range from 26-60 mm^2^ depending on the lamina^15,59,60^ so we scaled the average (x4) to 169 mm^2^ for humans. **(A)** Two very different-sized brushes (4 mm, orange; 16 mm, yellow) moved the same distance across the skin (16 mm left to right) can evoke similar global output neuron firing (pain intensity) if each stimulus only activates one LO neuron (green). Although different sizes, the two brushes evoke similar output firing because broader stimuli, by activating more of the inhibitory RF surround, engage more inhibition, which is reduced but not abolished by chloride dysregulation (dashed lines, e.g. in delay-spiking neuron: FR at 49 mm^2^ = FR at 306 mm^2^). **(B)** The same brushes as in A (4 and 16 mm), but moved for 52 mm, activate all four LO neurons and cause an increase in the firing rate of the global output neuron. As a result, both stimuli result in more intense pain than the stimulus design in A. **(C)** A larger brush size of 32 mm (which covers two LO neuron RFs) moved the same distance (16 mm) as the 4 mm brush results in more pain than the smaller brush.

**Figure S7.**
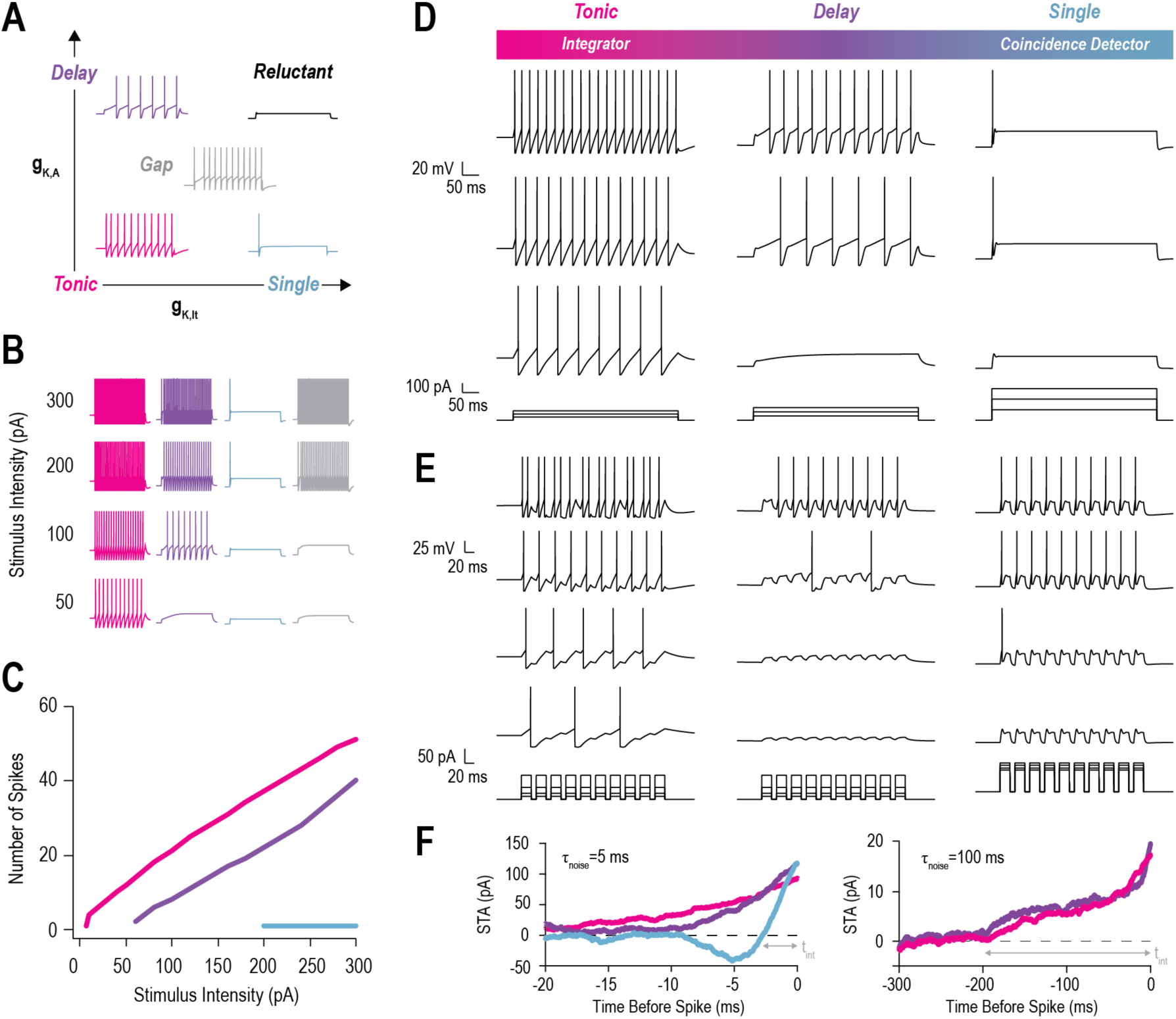
Models reproduce the operating mode of spinal neuron types. **(A)** Typical spiking patterns – single- (blue), delayed- (purple), gap- (grey) or reluctant- (black) spiking – observed in spinal neurons when injected with sustained current are reproduced by adding a transient K^+^ conductance (g_K,A_) and/or low threshold K^+^ conductance (g_K,lt_) to a basic tonic spiking AdEx model. **(B)** Injecting current steps of increasing intensity reproduces the spiking responses. **(C)** Models reproduce experimental F-I curves and capture neuron operating mode. **(D)** Tonic-spiking neurons (integrators) fire repetitively to sustained depolarization, delayed-spiking neurons respond with a short delay, and single-spiking neurons (coincidence detectors) respond with 1-2 spikes only at stimulus onset. Tonic-spiking neurons are the most sensitive to input (i.e. require the least total excitation to reach spike threshold). **(E)** Tonic-spiking neurons temporally summate input during pulse trains, delayed-spiking neurons only summate at higher amplitudes, and single-spiking neurons do not temporally summate input. The ability of tonic-spiking neurons to summate input is counterbalanced by a slow afterhyperpolarization (i.e. adaptation current) to prevent excessive firing, which can cause irregular spike timing. **(F)** Spike-triggered averages (STAs) for fast (τ_noise_ = 5 ms) or slow (τ_noise_ = 100 ms) current noise generated with an Ornstein-Uhlenbeck process (see Methods). Typical of their operating modes^19,20^, single-spiking neurons exhibited a biphasic STA and short integration time window (t_int_ ∼ 3 ms) for fast noise, whereas tonic- and delayed-spiking neurons exhibited monophasic STAs with long t_int_ (> 15 ms). Only tonic- and delayed-spiking neurons responded to slow noise, with both exhibiting a monophasic STA with t_int_ ∼ 200 ms. Single-spiking neurons did not respond to slow noise because they require fast fluctuation input to reach spike threshold^19,20^.

**Figure S8.**
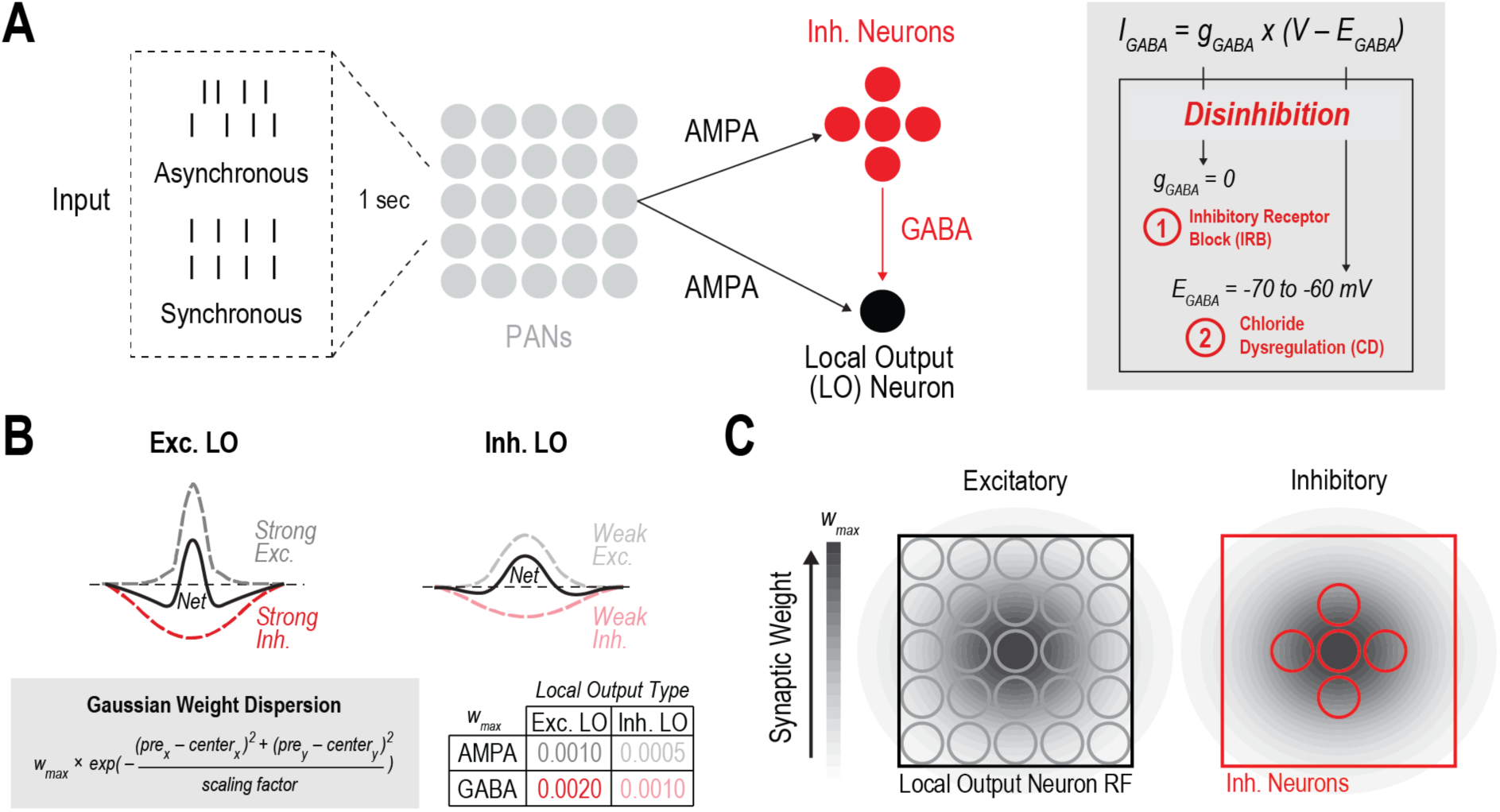
SDH circuit model. **(A)** PANs form AMPA synapses with inhibitory (Inh.) neurons and LO neurons. Touch input was modeled as sustained pressure or vibration, which causes asynchronous or synchronous firing in PANs. The breadth of the stimulus is parameterized as the number of PANs activated, which is set as a square (see Fig. 2C). For touch stimuli, all PANs within the square are uniformly activated, which is unlike the spatially disordered PAN activation caused by electrical stimulation (see Fig. 4). A touch stimulus can also move across PANs (see Fig. 3). Inhibitory neurons form GABA synapses on LO neurons. All inhibitory neurons are modeled as tonic-spiking. LO neurons were modeled with different spiking patterns but are delay-spiking if not otherwise mentioned. Disinhibition is simulated as inhibitory receptor block (IRB) or chloride dysregulation (CD). **(B)** Excitatory and inhibitory LO neurons have differential E-I balance based on experiments^15^. Excitatory LO neurons receive strong excitation (dark gray dashed) balanced by strong inhibition (red dashed), whereas inhibitory LO neurons receive weaker excitation (light gray dashed) balanced by weak inhibition (pink dashed). To capture this E-I balance, synaptic weights are spatially dispersed using a Gaussian function with a maximum weight (w_max_). The value of w_max_ changes varies with LO neuron type (Exc. LO or Inh. LO) and synapse type (AMPA or GABA). **(C)** Depiction of synaptic weight dispersion over space and scaling by w_max_.

**Table 1.**
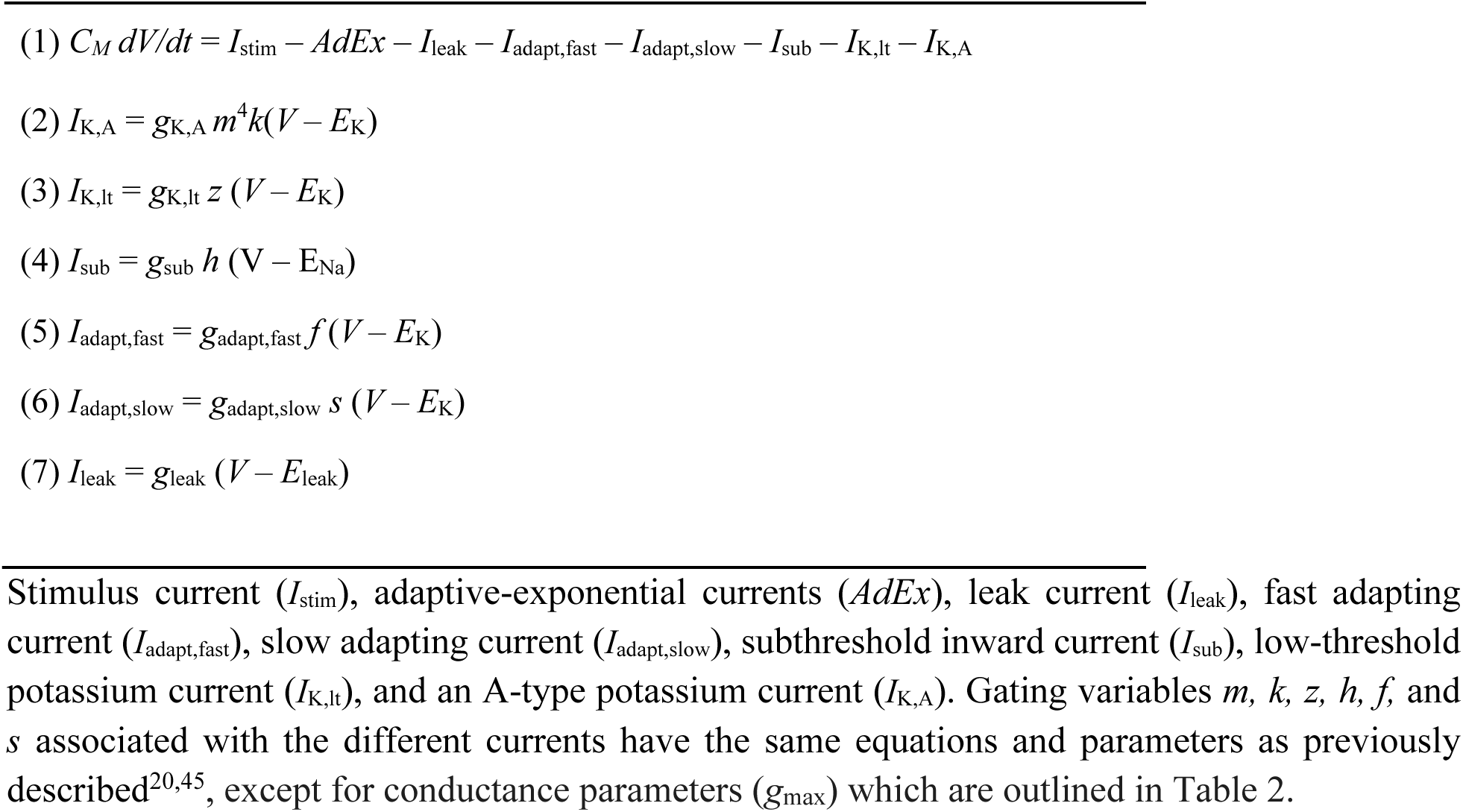
Equations for spinal neuron models.

**Table 2.** Parameters for spinal neuron models.

| Parameter | Tonic-spiking | Delayed-spiking | Single-spiking |
| --- | --- | --- | --- |
| $g_{leak}$ | 0.07 | 0.05 | 0.05 |
| $g_{adapt-fast}$ | 40 | 100 | 40 |
| $g_{adapt-slow}$ | 40 | 0 | 40 |
| $g_{sub}$ | 0.055 | 0 | 0 |
| $g_{K,A}$ | 0 | 2 | 0 |
| $g_{K,lt}$ | 0 | 0 | 15 |
| D | 30 | 30 | 25 |
| L | 28 | 28 | 23 |

Maximal conductance densities ( $g_{max}$ ) are in mS/cm<sup>2</sup>. Soma diameter (D) and length (L) are in $\mu$ m.

